# Inhibition of Mul1-mediated ubiquitination promotes mitochondria-associated translation

**DOI:** 10.1101/2021.07.28.454107

**Authors:** Yuan Gao, Maria Dafne Cardamone, Julian Kwan, Joseph Orofino, Ryan Hekman, Shawn Lyons, Andrew Emili, Valentina Perissi

## Abstract

G-Protein Pathway Suppressor 2 (GPS2) was recently identified as an endogenous inhibitor of non-proteolytic ubiquitination mediated by the E2 ubiquitin-conjugating enzyme Ubc13. GPS2-mediated restriction of K63 ubiquitination is associated with the regulation of insulin signaling, inflammation and mitochondria-nuclear communication, however a detailed understanding of the targets of GPS2/Ubc13 activity is currently lacking, Here, we have dissected the GPS2-regulated K63 ubiquitome in mouse embryonic fibroblasts and human breast cancer cells, unexpectedly finding an enrichment for proteins involved in RNA binding and translation. Characterization of putative targets, including the RNA-binding protein PABPC1 and translation factor eiF3m, revealed a strategy for regulating the mitochondria-associated translation of selected mRNAs via Mul1-mediated ubiquitination. Our data indicate that removal of GPS2-mediated inhibition, either via genetic deletion or stress-induced nuclear translocation, promotes the ubiquitination of mitochondria-associated translation factors leading to increased expression of an adaptive antioxidant program. In light of GPS2 role in nuclear-mitochondria communication, these findings reveal an exquisite regulatory network for modulating mitochondrial gene expression through spatially coordinated transcription and translation.

## INTRODUCTION

Maintenance of mitochondria metabolism and energy homeostasis is guaranteed by a constant turnover of the mitochondrial network, including both generation of new mitochondria, remodeling of existing mitochondria and removal of damaged organelles^1^. Mitochondria biogenesis and rewiring of mitochondrial functions require coordinated gene expression from the mitochondrial and nuclear genomes^2–5^. This is achieved through energy-sensing signaling pathways impacting on the transcriptional network controlling the expression of nuclear-encoded mitochondrial genes, and via shuttling of transcription factors and cofactors between the two organelles^6–9^. In yeast, balanced expression of different OXPHOS subunits is also facilitated by synchronized control of mitochondrial and cytosolic translation^2,10–13^, suggesting that coordinated regulation of multiple levels of gene expression may be required for the maintenance of mitochondrial homeostasis. However, the molecular mechanisms promoting balanced mitochondrial gene expression and specific mRNA processing and translation at the level of single organelles in mammals are not yet fully understood. Recent studies have identified RNA-binding proteins responsible for recruiting selected mRNAs to the mitochondrial outer membrane surface^14–23^, providing some clues to the specificity of translation on mitochondria-associated ribosomes as compared to cytosolic ribosomes. However a complete understanding of the modes of regulation of mitochondria-associated translation is currently lacking.

Ubiquitination is a reversible protein post-translational modification that is achieved through the sequential actions of several classes of enzymes, including a ubiquitin (Ub)-activating enzyme (E1), a Ub-conjugating enzyme (E2), and a Ub ligase (E3), that work together to mediate the attachment of one or multiple ubiquitin moieties to lysine residues on target proteins^24,25^. In the case of poly-ubiquitination, chains of different topology can either promote protein degradation or serve, as in the case of other post-translational modifications, to influence protein functions and interactions^26,27^. Among others, K48-linked ubiquitin chains are best known as markers for proteasomal degradation, whereas K63-linked ubiquitin chains are non-proteolytic^28,29^ and are associated with immune signaling, DNA damage repair, protein sorting, and translation^30–37^. Moreover, more recent studies point to a broader role in regulating cellular metabolism and adaptive responses to stress^38–43^. Notably, K63-linked ubiquitin mediates translational regulation in yeast exposed to oxidative stress^39,42^.

We recently identified GPS2 as an inhibitor of K63 ubiquitination synthesized by the E2 conjugating enzyme Ubc13^44^. GPS2-mediated inhibition of K63 ubiquitination is essential for promoting several coordinated functions across the cell, including the transcriptional regulation of mitochondrial genes in the nucleus and activating insulin signaling and pro-inflammatory pathways in the cytosol^9,44–47^. These complementary activities across different subcellular compartments are facilitated by GPS2 translocation between organelles, making GPS2 an excellent candidate for mediating the coordination of nuclear and extranuclear processes contributing to the remodeling of cellular metabolism. Regulated mitochondria-to-nucleus shuttling of GPS2, for example, controls mitochondria biogenesis via transcriptional activation of nuclear-encoded mitochondrial genes^9^. Acting as a mediator of mitochondria retrograde signaling, GPS2 is also required for licensing the expression of stress response genes upon mitochondria depolarization^9^. Upon translocation to the nucleus, GPS2-mediated inhibition of K63 ubiquitination impacts the remodeling of the chromatin environment of target gene promoters by stabilizing histone demethylases^9^. At the same time, its removal from the outer mitochondrial membrane may release the ubiquitination of other targets, which are currently unknown. Here, we have profiled the GPS2-regulated ubiquitome and dissected the role of GPS2-regulated K63 ubiquitination in modulating mitochondria-associated protein translation. Our results point to a novel regulatory strategy whereby GPS2 provides a unifying strategy for coordinating the transcriptional and translational control of mitochondrial gene expression by inhibiting K63 ubiquitination of key targets by the mitochondrial ubiquitin ligase Mul1.

## RESULTS

### Remodeling of the mitochondrial proteome in the absence of GPS2

Our previous work has identified GPS2 as a regulator of mitochondrial gene expression, through transcriptional control of nuclear-encoded mitochondrial (neMITO) genes, and as a mediator of mitochondria-to-nucleus retrograde signaling^9^. To further dissect the role of GPS2 in regulating mitochondrial functions, we profiled the mitochondrial proteome and transcriptome of immortalized mouse embryonic stem cells (MEFs) derived from either GPS2^fl/fl^/Ubc^ERT2^-Cre^+^ or GPS2^fl/fl^/Ubc^ERT2^-Cre^−^ mice. Upon Tamoxifen treatment, GPS2 deletion is induced in Ubc^ERT2^-Cre^+^ cells (from here on called GPS2-KO), whereas Ubc^ERT2^-Cre^−^ (from here on called WT) serve as negative control (**Supplemental Figure S1A)**. In agreement with previous studies in other cell models, GPS2-KO MEFs present with severely reduced mitochondrial content compared to WT cells (**Supplement Figure S1B**).

To achieve a comprehensive characterization of the changes associated with GPS2 deletion, we first profiled mitochondrial extracts from WT and KO MEFs using quantitative tandem mass tag (TMT) labeling followed by quantification via mass spectrometry (LC-MS/MS). We identified 891 differentially expressed proteins across three replicates (**Supplementary Table 1**, P-value <0.05 and LogFC>0.25). In accord with our previous findings in adipocytes and breast cancer cells^44,46,47^, upregulated proteins were enriched for pathways previously found to be regulated by cytosolic GPS2, including EGFR/Insulin signaling, inflammatory responses, and TNFalpha/MAPK signaling (**Supplemental Table 1** and **Figure S1C**). Downregulated proteins, instead, spanned various mitochondrial functions, including Electron Transport Chain (ETC), fatty acid oxidation, TCA cycle, and amino acid metabolism (**Supplemental Table 1** and **Figure S1C**). Accordingly, filtered analysis of intrinsic mitochondrial proteins, based on the latest MitoCarta 3.0 database, indicated that the majority of differentially expressed mitochondrial proteins are downregulated in GPS2-KO cells, as expected based on previous studies (**Figure 1A**)^9^. However, we also identified a small subset of mitochondrial proteins upregulated in the absence of GPS2. Interestingly, the upregulated program was enriched in proteins involved in antioxidant response and programmed cell death (**Figure 1A**), which suggests adaptive changes in response to the stress of GPS2 deletion. Upregulation of anti-oxidative protective proteins SOD2 and PRDX2, fatty acid synthase FASN, and downregulation of pyruvate dehydrogenase PDHA and OXPHOS proteins was confirmed by western blotting of mitochondrial extracts from GPS2 WT and KO MEFs (**Figure 1B**).

**Figure 1.**
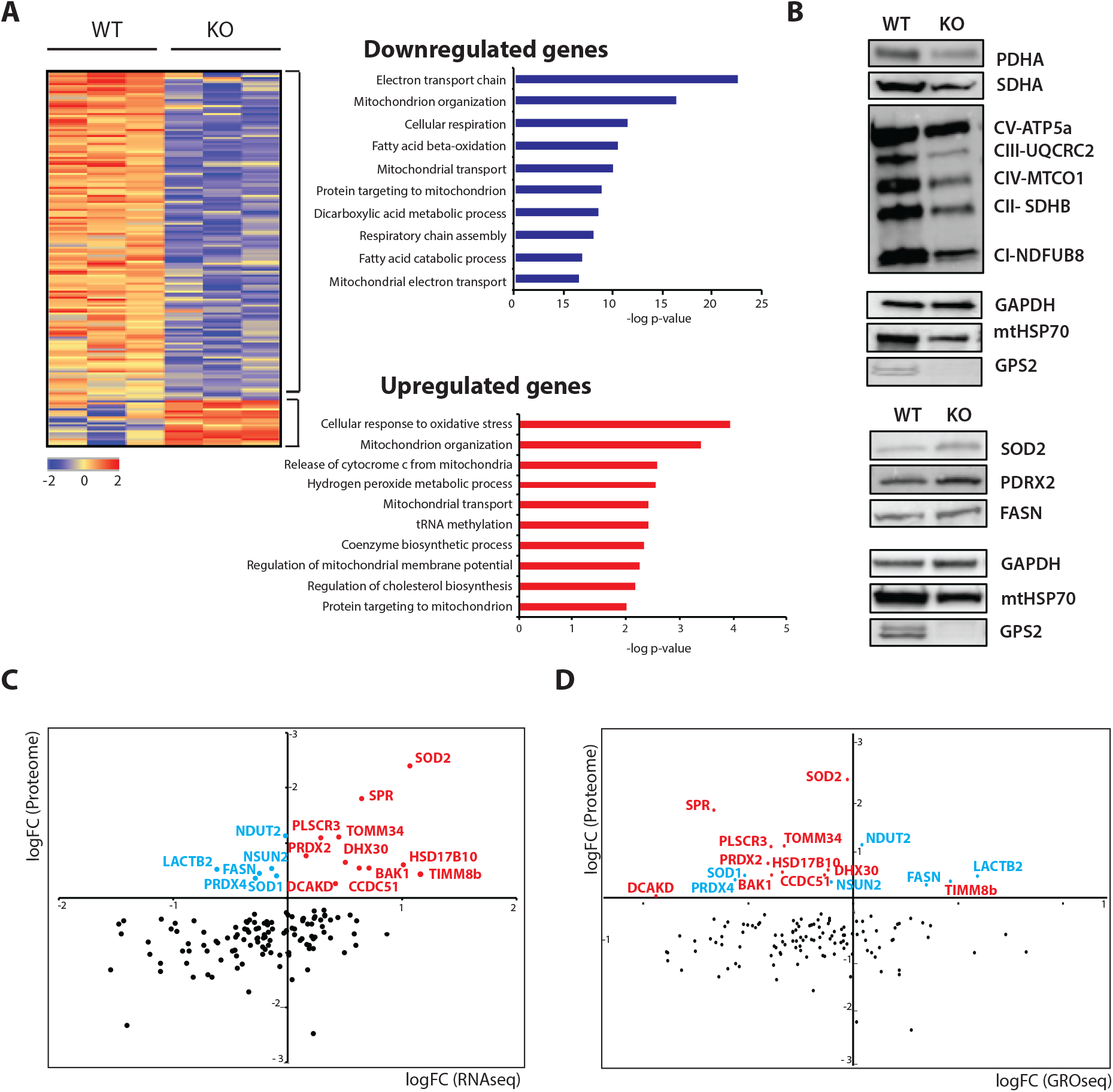
Mitochondrial proteome of GPS2 WT and KO MEFs. A, Heatmap showing 137 significantly changed mitochondrial proteins induced by GPS2-deficiency from GPS2 WT and KO MEFs. Enriched biological process GO terms of significantly downregulated (blue, upper panels) or upregulated (red, down panels) in response to GPS2 deficiency. B, Western blot analysis of SOD2, PDRX2, and FASN normalized to GAPDH in mitochondrial extracts from WT versus GPS2 KO MEFs. C&D, Distribution of 137 significantly changed mitochondrial proteins based on LC-MS/MS analysis and RNA-seq (C) or previously generated GRO-seq in 3T3-L1 cells^9^. Red, upregulated proteins identified through RNA-seq and proteomics; Blue, upregulated proteins identified through proteomics, but downregulated in RNA-seq; Black, downregulated proteins identified through proteomics.

To characterize the molecular mechanisms underlying the adaptive regulation of mitochondrial gene expression in the absence of GPS2, we profiled mitochondria-associated RNA from WT and GPS2-KO MEFs by RNA-seq. Among the differentially expressed genes (DEGs) (**Supplementary Table 2**), we identified 316 mitochondrial genes consistently regulated across three independent replicates, including 138 upregulated and 178 downregulated genes. Overlay of the RNA-seq results over the proteomics data confirmed that the downregulation of mitochondrial proteins observed in GPS2-KO cells is largely associated with reduced mRNA levels (**Figure 1C**), as expected based on GPS2 acting as a required transcriptional cofactor for the activation of neMITO genes^9^. Indeed, overlay of the MEFs proteomics data over GRO-seq data previously generated in 3T3-L1 cells with acute GPS2 downregulation further indicates that protein downregulation likely results from defective transcription (**Figure 1D**)^9^. Mitochondrial proteins upregulated in GPS2-KO cells instead included genes presenting both increased (59%, depicted in red in **Figure 1C**) or decreased (35%, depicted in blue in **Figure 1C**) mRNA abundance in KO versus WT cells. However, a comparison of protein and nascent RNA expression suggests that the transcription of the majority of proteins differentially regulated in MEFs, including both up- and down-regulated genes, is impaired in the absence of GPS2 (**Figure 1D**). This suggests that multiple levels of regulation may be contributing to the remodeling of the mitochondrial proteome of GPS2-KO cells, including both transcriptional repression and post-transcriptional regulation of mRNA stability and/or protein synthesis, with the outcome for different gene/protein sets depending on the overlay of these complementary regulatory strategies.

### GPS2 restricts K63 ubiquitination of mitochondrial proteins

To further our understanding of the adaptive rewiring of the mitochondrial proteome in GPS2-null cells, we decided to investigate the mechanism/s responsible for post-transcriptional regulation of mitochondrial gene expression in the absence of GPS2. Because our previous work indicates that GPS2 exerts its functions across different subcellular compartments via inhibition of the ubiquitin-conjugating enzyme Ubc13^9,44–48^, we focused on K63 ubiquitination as a possible regulatory mechanism. First, we asked whether loss of GPS2 leads to unrestricted K63 ubiquitination activity on mitochondria. Indeed, we observed increased accumulation of K63 ubiquitin chains in mitochondrial extract from GPS2 KO cells compared to the WT line by western blot analysis (**Figure 2A**). To identify putative targets of GPS2-mediated regulation, we then profiled the K63 ubiquitome of GPS2 WT and KO MEFs by LC-MS/MS. Relative quantification was performed by stable isotope labeling of amino acid in cell culture (SILAC) (**Figure 2B**), with K63 ubiquitinated proteins being enriched prior to mass spectrometry through selective binding to the high-affinity lysine-63-poly-ubiquitin binding domain Vx3K0^49^ (**Supplemental Figure 2A** and **2B**). This approach led to the identification of 73 candidate targets, among 230 differentially enriched proteins, for which the increase in the H/L ratio indicated increased ubiquitination and/or increased interaction with ubiquitinated targets in GPS2-KO cells (**Supplemental Figure 2C** and **Supplemental Table 3**). Unexpectedly, we found that putative targets of GPS2-mediated regulation were strongly enriched for factors involved in protein translation (**Figure 2C**), whereas no significant overlap with known targets of K63 ubiquitination by Parkin was observed, despite mitophagy markers being activated in GPS2-KO cells (Supplemental Figure 2D, 2E and **Supplemental Table 3**). These findings were confirmed when profiling was repeated using TMT labeling instead of SILAC (**Supplemental Figure 2D**). A similar enrichment in proteins involved in translation and RNA processing was also observed by profiling the K63 ubiquitome of MDA-MB231 breast cancer cells deleted of GPS2 (REF) (**Supplemental Figure 2E** and **Supplemental Table 3**). Moreover, the increased ubiquitination of representative translation factors RPS11 and RACK1 - both previously reported as direct targets of non-proteolytic ubiquitination (REFs) - was confirmed by IP/WB using both whole-cell extracts and fractionated mitochondrial extracts (**Figure 2D**). This indicates that lack of GPS2 promotes exacerbated ubiquitination of translation factors associated with mitochondria, thus suggesting that GPS2-mediated inhibition of K63 ubiquitination might regulate mitochondria-associated protein translation.

**Figure 2.**
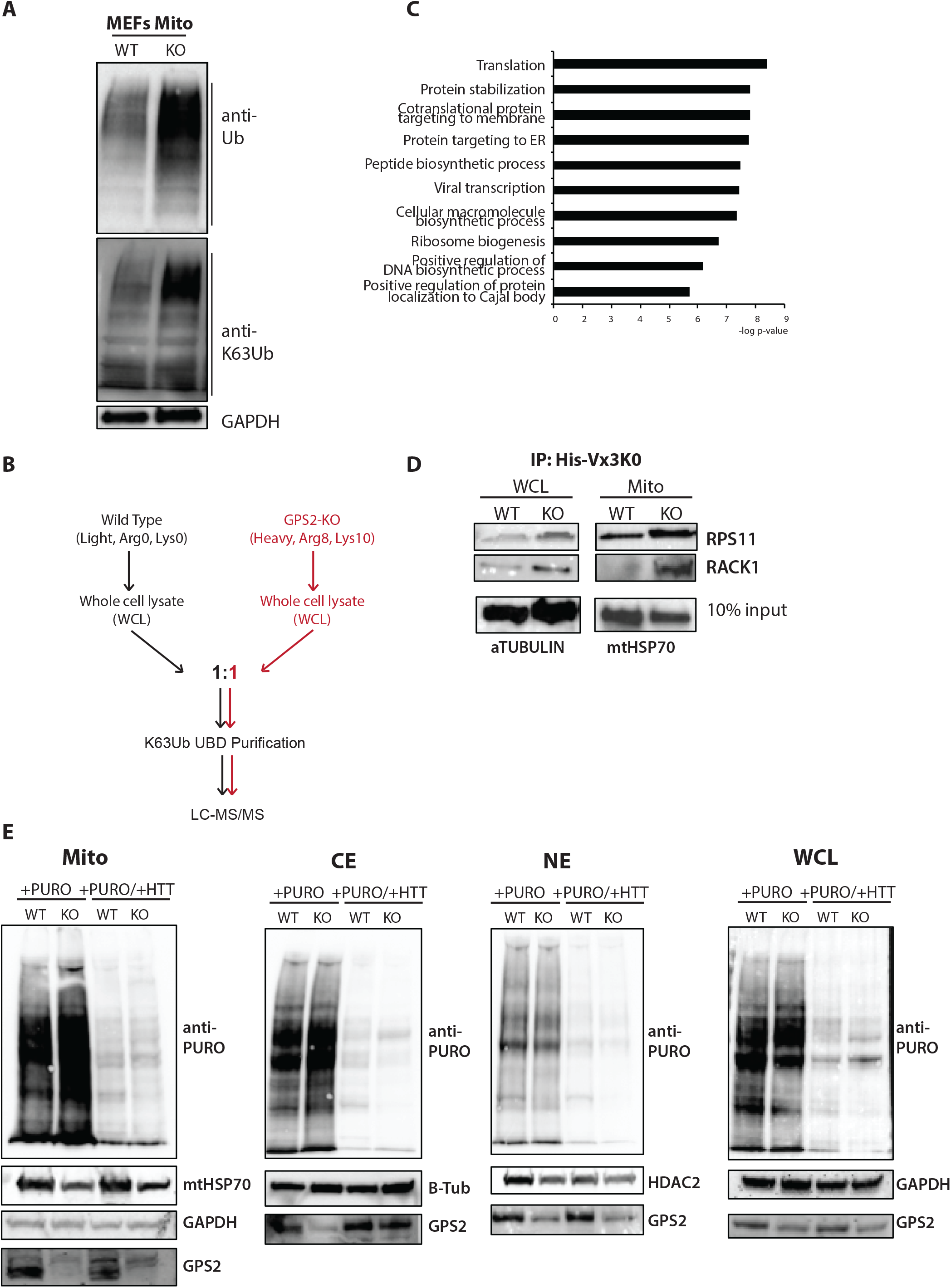
K63 ubiquitome profiling of GPS2 WT and KO MEFs by LC-MS/MS. A, Increased K63 ubiquitination in mitochondria extract in GPS2 KO compared to WT by western blot. B, SILAC-based K63 proteomic profiling of GPS2 WT and KO MEFs. C, Gene ontology analysis of 46 putative K63 ubiquitinated proteins whose ubiquitination is mediated by GPS2. D, Detection of K63 ubiquitination of RPS11 and RACK1 purified by specific K63-ubiquitin binding domain Vx3K0 by IP/WB using both whole-cell extracts and fractionated mitochondrial extracts in GPS2 WT and KO MEFs. E, Puromycin labeling of *de novo* protein synthesis on different fractionated extracts pretreated w/o homoharringtonine (HHT). ME, mitochondrial extract; CE, cytosol extract; NE, nuclear extract; WCL, whole cell extract. Cells were treated with HHT for 10 min prior to treatment with 0.9mM puromycin for additional 5 min. Anti-GAPDH/B-Tub/HDAC2 blot was used as loading control for different fractions.

To investigate whether protein translation is regulated in absence of GPS2, we compared the rate of protein translation in WT and KO cells by monitoring the incorporation of puromycin into newly synthesized proteins across subcellular compartments. A significant increase in puromycin incorporation was observed in mitochondria from GPS2-KO cells compared to WT (**Figure 2E**). Pre-treatment of cells with the translation initiation inhibitor homoharringtonine (HHT)^50^ reduced puromycin incorporation in both WT and KO mitochondria, indicative of active translation. Remarkably, the increase in puromycin incorporation was specific to the mitochondrial compartment, with no significant changes observed in nuclear and cytosolic extracts (**Figure 2E**). Together, these results indicate that the rate of protein translation in mitochondrial extracts is specifically upregulated in absence of GPS2. Based on previous studies indicating that this approach allows for selective measurement of translation of nuclear-encoded mitochondrial proteins occurring on the outer mitochondrial membrane (OMM) as compared to the translation of mitochondria-encoded genes^51^, we concluded that mitochondria-associated translation of neMITO genes increases in absence of GPS2. This suggests a localized regulatory strategy impacting mRNAs translated by ribosomes docked on the mitochondria rather than free cytosolic ribosomes. In accord with this hypothesis, we observed that transcripts encoding for more than half of the upregulated proteins in GPS2-KO cells are recruited to the outer mitochondrial membrane for import-coupled translation via interaction with the AKAP1/MDI/LARP complex^18,52^(**Supplementary Table 1**). Conversely, no significant overlap was observed between upregulated proteins in the absence of GPS2 and transcripts binding to CluH, an RNA binding protein responsible for the assembly of cytosolic granules promoting the translation of a variety of mitochondrial proteins^14,15,53^, including several showing reduced transcription in the absence of GPS2 (**Supplementary Table 1**). Together, these comparative *in silico* analyses suggest that enhanced translation of mitochondrial proteins in the absence of GPS2 is specific for mitochondrial transcripts translated by mitochondria-associated ribosomes.

### PABPC1 ubiquitination by Mul1

To gain further mechanistic insights into the regulation of mitochondria protein translation via K63 ubiquitination, we focused on the poly-A binding protein PABPC1. PABPC1 is not only a known target of ubiquitination^54^ but also the single putative target of GPS2 regulation identified across different multiple experimental setups (SILAC and TMT labeling) and different cell types (MEFs and MDA-MB-231) (**Supplemental Figure 3A**). Two complementary IP/WB approaches confirmed increased K63 ubiquitination of PABPC1 in the mitochondria of GPS2-KO cells. In the first assay, we immunoprecipitated K63Ub-containing proteins by binding to the Vx3KO trap and visualized PABPC1 by WB (**Figure 3A**). In the second, we immunoprecipitated PABPC1 and visualized associated K63Ub chains by WB (**Figure 3B**). As expected, the increase in high molecular weight (HMW) Ub chains observed in GPS2-KO cells was reversed by inhibiting the activity of ubiquitin-conjugating enzyme Ubc13 with the chemical inhibitor NSC697923 (REF)(**Figure 3B**). A Ubc13-dependent increase in PABPC1 ubiquitination was also observed upon depolarization of mitochondria by FCCP, a condition mimicking GPS2 downregulation through its relocalization from mitochondria to the nucleus^9^(**Figure 3C**). Unexpectedly, this experiment revealed that FCCP-induced ubiquitination of PABPC1 depended on the mitochondria specific E3 ligase MUL1 rather than MKRN1, which had been previously associated with PABPC1 ubiquitination in the cytosol (**Figure 3C**). Ubiquitination of PABPC1 by MUL1 and Ubc13, as recapitulated *in vitro* using bacterially expressed recombinant proteins, was significantly reduced upon mutagenesis of the previously identified ubiquitination sites (K78, K188, K284, and K512) (**Figure 3D**) and significantly inhibited by GPS2 (**Figure 3E**). Together these results indicate that PABPC1 can be locally regulated through different ubiquitination machineries and that GPS2 is specifically responsible for preventing its ubiquitination by Mul1 on mitochondria.

**Figure 3.**
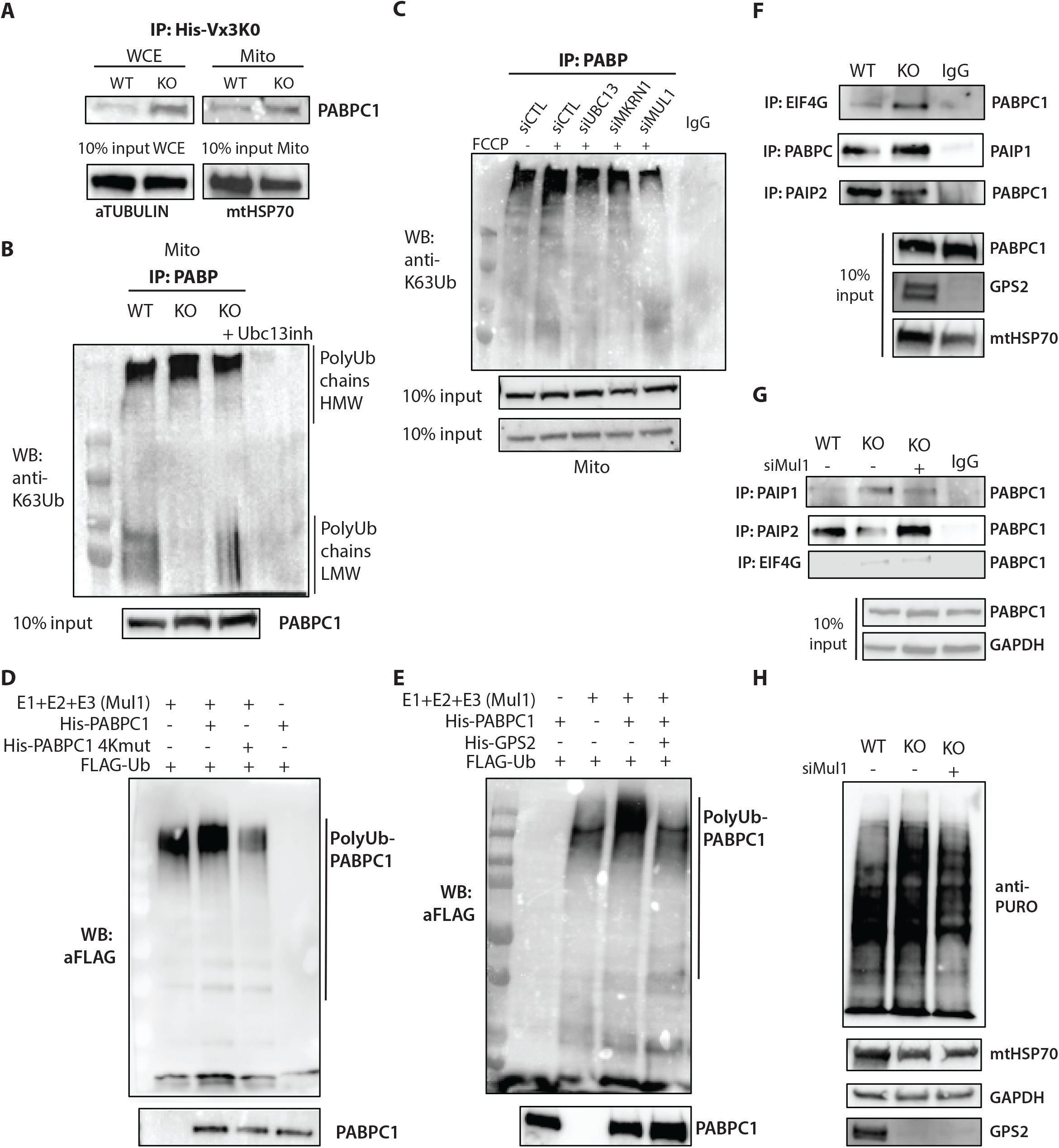
PABPC1 ubiquitination is mediated by Mul1 on mitochondria. A, Detection of K63 ubiquitination of PABPC1 purified by specific K63-ubiquitin binding domain Vx3K0 by IP/WB using both whole cell extracts and fractionated mitochondrial extracts in GPS2 WT and KO MEFs. B, Detection of K63 ubiquitination by IP/WB of PABPC1 in mitochondria extracts in GPS2 WT, KO and KO treatment with UBC13 inhibitor MEFs. C, FCCP-induced ubiquitination of PABPC1 is mediated by mitochondria specific E3 ligase MUL1. D, PABPC1 is polyubiquitinated by the E3 ubiquitin ligase MUL1 by *in vitro* ubiquitination assay. E, Polyubiquitination on PABPC1 mediated by E3 ubiquitina ligase MUL1 is significantly inhibited by GPS2 in *in vitro* ubiquitination assay. F, WB analysis of binding of PABPC1 with EIF4G, PAIP1 and PAIP2 on mitochondrial extracts from GPS2 WT and KO MEFs. G, WB analysis of binding of PABPC1 with PAIP1 and PAIP2 on mitochondrial extracts from GPS2 WT, KO and KO knockdown MUL1 (siMul1) MEFs. H, Puromycin labeling of *de novo* protein synthesis on mitochondrial extracts from GPS2 WT, KO and KO knockdown MUL1 (siMul1) MEFs. Anti-GAPDH blot was used as loading control.

PABPC1 is an RNA-binding protein and a key component of the translation machinery. PABPC1 concomitant interactions with the polyA-containing 3’UTR and 5’ Cap-bound components of the translation initiation complex promote protein translation through closed-loop mRNP formation^55,56^. To unravel whether loss of GPS2 may be impacting on mitochondria-associated translation through aberrant PABPC1 ubiquitination, we first asked whether PABPC1 interaction with the translation initiation complex is altered in the absence of GPS2. Co-immunoprecipation experiments indicated that PABPC1 interaction with the translation initiation factor eIF4G and translational activator PABP-interacting protein 1 (PAIP1) is more robust in mitochondria from GPS2-KO cell than their WT counterparts (**Figure 3F)**. In contrast, interaction with the translational repressor PABP-interacting protein 2 (PAIP2), which competes with PAIP1 for interaction with PABP^57^, is significantly decreased (**Figure 3F)**. These changes are rescued by transient downregulation of the E3 ubiquitin ligase MUL1 (**Figure 3G**), indicating that aberrant ubiquitination of MUL1 target/s in the absence of GPS2-mediated inhibition of Ubc13 activity is indeed a contributing factor to the observed phenotype. This conclusion was further confirmed by rescue of enhanced protein translation rate in GPS2-KO MEF cells through MUL1 downregulation by siRNA, as measured by mitochondrial puromycilation assay (**Figure 3H**).

While we have primarily focused on PABPC1 due to its striking identification across multiple datasets, we also considered the possibility that ubiquitination of additional factors may be contributing to regulate mitochondria-associated translation. GPS2-regulated K63 ubiquitome included a number or putative targets involved in RNA processing and translation. Among them, eukaryotic translation initiation factor 3, subunit M (eiF3m), was also confirmed as a direct target of GPS2/Ubc13/Mul1-mediated regulation (**Supplemental Figure 3B**). Our results together indicate that GPS2 regulates mitochondria-associated translation by preventing aberrant MUL1-mediated ubiquitination of translation factors like PABPC1 and eIF3m on the outer mitochondrial membrane.

### K63ub-mediated regulation of mitochondria-associated translation in stress response

Our previous studies indicate that, in conditions of mitochondrial depolarization, GPS2 translocates to the nucleus for regulating the transcription of stress response genes^9^. This suggested that removal of GPS2 by translocation could have the indirect effect of releasing Mul1 activity and promoting mitochondrial protein translation as part of the adaptive stress response. To investigate this possibility, we first assessed the ubiquitination of PABPC1 after depolarization with FCCP for 10’ or 30’. As expected, concomitant to the translocation of GPS2 from mitochondria to the nucleus (**Figure 4A**), we observed a significant increase in mitochondria-associated K63 poly-ubiquitination of PABPC1 (**Figure 4B** and **4C**). Under these conditions, PABPC1 interaction with the translation initiation complex is strengthened (**Figure 4D**). This correlates with the activation of a gene expression program that, in part, mirrors that identified in response to GPS2 deletion (**Figure 4E, bottom cluster**), as validated by western blotting for antioxidant proteins SOD2 (**Figure 4F**). At the same time, activation of a large part of the FCCP-induced program is impaired in GPS2-KO cells, likely due to reduced transcription in the absence of nuclear GPS2.

**Figure 4.**
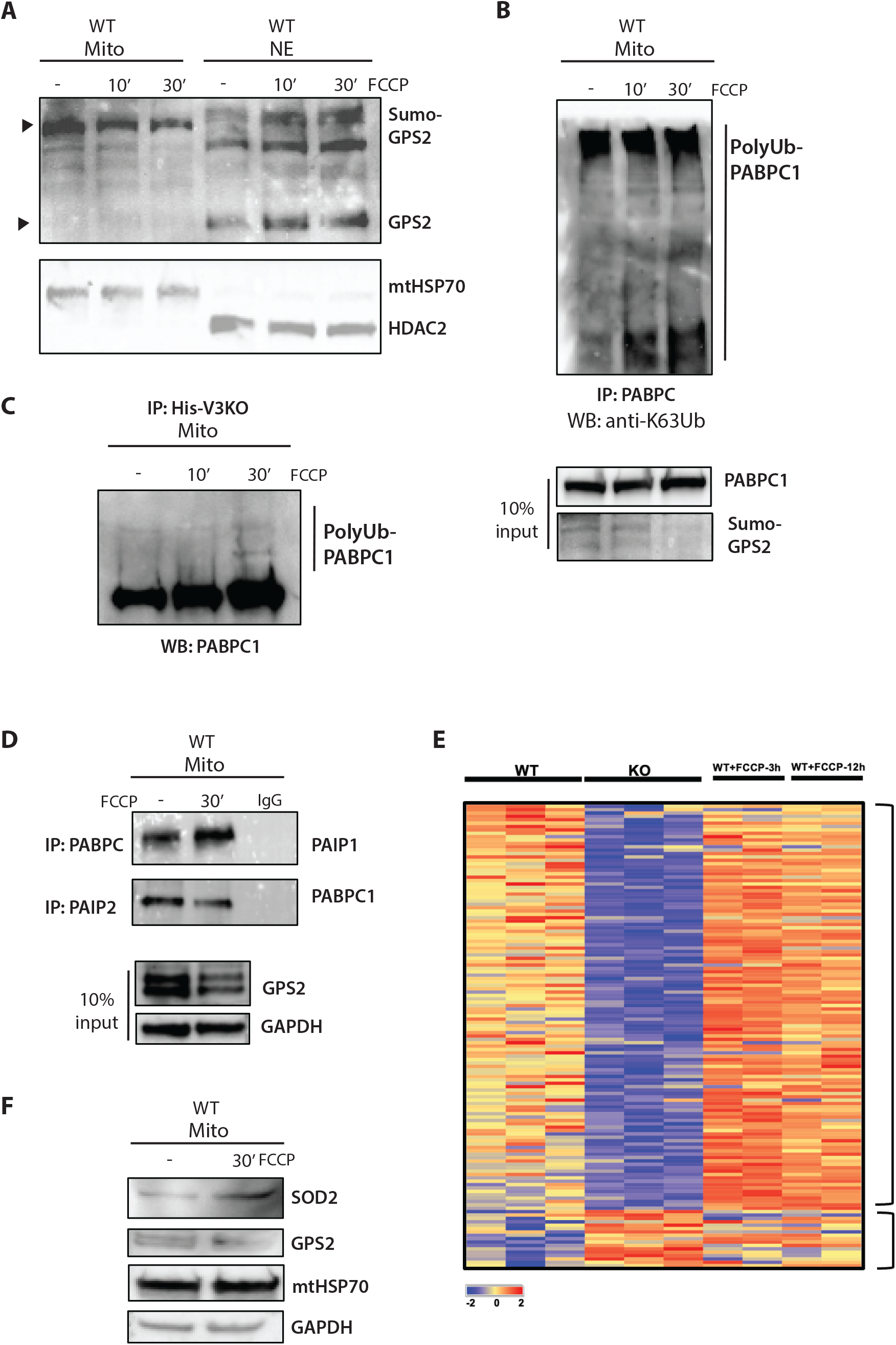
K63 ubiquitination-mediated regulation of mitochondrial-associated translation in stress response. A, WB of fractionated extracts showing mitochondria-to-nucleus translocation of GPS2 in MEFs cells upon FCCP treatment. B, Detection of K63 ubiquitination by IP/WB of PABPC1 in mitochondria extracts in MEFs cells upon FCCP treatment. C, Detection of K63 ubiquitination of PABPC1 purified by specific K63-ubiquitin binding domain Vx3K0 by IP/WB in mitochondria extracts in MEFs cells upon FCCP treatment. D, WB analysis of binding of PABPC1 with PAIP1 and PAIP2 in mitochondria extracts in MEFs cells upon FCCP treatment. E, Heatmap showing significantly changed mitochondrial proteins in GPS2 WT, KO MEFS and WT cells treated with FCCP for 3h and 12h. F, Western blot analysis of SOD2 normalized to GAPDH in mitochondrial extracts in MEFs cells upon FCCP treatment.

## DISCUSSION

Previous studies have characterized GPS2 as a mediator of mitochondria retrograde signaling, a transcriptional cofactor involved in mediating both gene repression and activation, and an inhibitor of non-proteolytic K63 ubiquitination. Together, these complementary functions ensure a proper adaptive response to mitochondrial stress and regulate mitochondria biogenesis during differentiation, with GSP2 translocating from mitochondria to nucleus to regulate nuclear-encoded mitochondrial genes transcription via stabilization of chromatin remodeling enzymes^9^. A key implication of these findings is that restricted K63 ubiquitination in the nucleus indirectly regulates mitochondrial functions through the expression of stress response and nuclear-encoded mitochondrial genes. However, it remains unclear whether GPS2 also contributes to the regulation of mitochondria homeostasis in a direct manner, possibly by modulating the ubiquitination of mitochondrial proteins. To address this question, we have profiled the GPS2-regulated K63 ubiquitome in mouse embryonic fibroblasts and human breast cancer cells. To our surprise, we did not observe any significant overlap between the GPS2-regulated program and the mitochondrial proteins targeted by Parkin-mediated ubiquitination. This suggests that stress-induced translocation of GPS2 is unlikely to contribute to removing defective mitochondria via mitophagy. Instead, GPS2-mediated inhibition of K63 ubiquitination prevalently affects proteins involved in RNA processing and translation. In particular, our results indicate that GPS2-mediated inhibition of K63 ubiquitination contributes to regulating the mitochondria-associated translation of a specific of nuclear-encoded mitochondrial genes. Together with the previous characterization of nuclear GPS2 as an essential cofactor for the expression of nuclear-encoded mitochondrial genes, these findings indicate that GPS2-mediated inhibition of K63 ubiquitination represents a unifying strategy for modulating the expression of mitochondrial proteins through coordinated transcription and translation events.

Mitochondrial proteins regulated through this strategy are enriched for factors involved in the antioxidant response, which is consistent with both the role played by GPS2 in mediating the mitochondrial stress response and the role played by K63 ubiquitination in regulating ribosomal activity in yeast (REF). Recent studies have identified RNA binding proteins, such as Puf3p in yeast, Clu and MDI/Larp in flies, CluH and dAKAP1/LARP4 in mammalian cells, that are involved in the selective regulation of different subsets of nuclear-encoded mitochondrial proteins^14,17,18,52,58–62^. Mitochondrial proteins upregulated in GPS2-KO cells present a significant overlap with the AKAP1/MDI/LARP complex targets, which is responsible for recruiting mRNAs to the outer mitochondrial membrane for import-coupled translation. In contrast, no overlap was observed with transcripts binding to CLUH, an RNA binding protein responsible for the assembly of cytosolic granules promoting the cytosolic synthesis of various mitochondrial proteins. These results suggest that different sets of nuclear-encoded mitochondrial genes might be regulated through independent strategies, with GPS2-controlled K63 ubiquitination being restricted to the local regulation of mitochondria-associated translation. This separation is likely to be critical for ensuring the maintenance of mitochondrial and cellular homeostasis.

Mechanistically, our results indicate that GPS2 regulates mitochondrial-associated translation by inhibiting the ubiquitination of the RNA binding protein PABPC1 and other translation factors by the mitochondrial E3 ubiquitin ligase Mul1. To our surprise, our data point to the mitochondria-specific Mul1 as the E3 ligase regulating PABPC1 ubiquitination both *in vivo* and *in vitro*, rather than MKRN1, which had been previously associated with the ubiquitination of PABPC1 in HEK293T^54^. Mul1 is a dual function E3 ligase promoting either the ubiquitination or SUMOylation of mitochondrial targets^63,64^. Previous studies have described an important role for Mul1 in regulating a variety of physiological and pathological processes, including mitochondrial dynamics, cell growth, apoptosis, and mitophagy^65–69^. Our study adds to this body of work by showing that Mul1-mediated ubiquitination regulates the activity of translation factors localized to the OMM and restricts the local translation of a subset of nuclear-encoded mitochondrial proteins. One intriguing aspect of Mul1 involvement in regulating mitochondria-associated translation is that it is uniquely suited for playing a key role in the mitochondrial stress response (MSR). Because of its spatial organization, Mul1 can in fact sense mitochondrial stress in the intermembrane mitochondrial space and respond through the ubiquitination of specific substrates on the outer mitochondrial surface^68–70^. Our data support this possibility as we observed increased ubiquitination of PABPC1 upon mitochondrial depolarization, concomitant to GPS2 retrograde translocation to the nucleus.

The relationship between GPS2 and Mul1 in mitochondria appears quite complex. Mul1 was previously shown to regulate the sumoylation of GPS2, as required for regulating GPS2 intracellular localization and translocation in stress conditions^9^. At the same time, results presented here indicate that GPS2 inhibits Ubc13/Mul1-mediated ubiquitination of PABPC1, likely through inhibition of Ubc13 activity. Together, these observations suggest a complex regulatory strategy for modulating mitochondrial adaptation to stress which includes: 1) Mul1-mediated sumoylation of GPS2 is removed by SENP1; 2) desumoylation favors GPS2 translocation to the nucleus where it promotes the transcription of stress response and nuclear-encoded mitochondrial genes; 3) at the same time the absence of GPS2 in mitochondria licenses Mul1/Ubc13 activity on mitochondria-associated translation factors, in turn enhancing the translation of antioxidant proteins to promote adaptation to the mitochondrial stress.

While we have mainly focused on PABPC1 as a key target of GPS2-regulated ubiquitination in this study, our proteomics data suggest that PABPC1 is not the only target relevant to the regulation of mitochondrial-associated translation. Ribosomes are known to be regulated by non-proteolytic ubiquitination as well, and ribosomal subunit RPS11 and scaffold factor RACK1^71–73^. In addition, we identified Eif3m, a translation initiation factor, as a target of GPS2-regulated K63 ubiquitination and GPS2. These results suggest that regulation of mitochondrial-associated translation is achieved by the concomitant ubiquitination of multiple factors working together.

Further studies will be required to dissect the specific sites of ubiquitination and the contribution of individual PTM events.

In conclusion, our work indicates that GPS2 plays an important role in mitochondrial regulation by inhibiting ubiquitin signaling. Together with the recent characterization of GPS2-mediated regulation of nuclear-encoded mitochondrial genes transcription, our results add to the significance of the GPS2 role in integrating multiple layers regulating cell growth, metabolism, and stress resistance.

## ACKNOWLEDGMENTS

We thank all past and present members of the Perissi Lab for their continuous support and encouragement. We are grateful to Drs. Christine Vogel and Gustavo Silva for their assistance with protocols for the enrichment of K63 ubiquitinated proteins. Genomic profiling experiments have been run by the Boston University Microarray & Sequencing Resource Core. This work was supported by NIH and Department of Defense grant awards as follows: NIGMS R01GM127625 to VP, DoD W81XWH-17-1-0048 to VP, R00GM124458 to SML

## Materials and Methods

### Reagents and antibodies

Anti-GPS2 antibody was generated in rabbit against a peptide representing aa 307-327. Commercial antibodies used were as follows: anti-ubiquitin (P4D1 clone, Cell Signaling Technology), anti-β-tubulin (TUB 2.1 clone, Sigma), anti-HDAC2 (ab16032, Abcam), anti-mtHSP70 (catalog no. MA3-028, Invitrogen), anti-K63 (catalog no. 05-1308, Millipore), anti-α-Tubulin (catalog no. T5168, Sigma), anti-GAPDH (catalog no. MA5-15738, Invitrogen), anti-PABPC1 (ab21060, Abcam), anti-RPS11(NBP2-22289, Novus Biologicals), anti-Rack1 (sc-17754, Santa Cruz Biotechnology), anti-Eif4G (15704-1-AP, Proteintech), anti-Paip1 (sc-365687, Santa Cruz Biotechnology), anti-Paip2 (15583-1-AP, proteintech), anti-Flag-HRP (catalog no. a8592, Sigma), anti-Puromycin (clone 12D10, Millipore), anti-PDHA (ab168379, Abcam), anti-Oxphos (ab110413, Abcam), anti-SOD2 (sc-137254, Santa Cruz Biotechnology). siRNAs against mouse GPS2, UBC13, MKRN1, MUL1, and UBC9 were purchased from Ambion. Nonspecific scrambled siRNA was included as negative controls in each experiment.

### Cell culture

Standard molecular cloning, cell culture, and cell transfection experiments were performed as described by J. Sambrook, D. W. Russell, Molecular Cloning: A Laboratory Manual (Cold Spring Harbor Laboratory Press, Cold Spring Harbor, N.Y., ed. 3rd, 2001). For cells transfection, jetPRIME was used following the manufacturer’s protocol (Polyplus transfection). Immunostaining was performed following standard protocols on cells fixed in 4% paraformaldehyde in PBS, using Alexa Fluor-conjugated secondary antibodies (Molecular probes).

Mouse embryonic fibroblasts (MEFs) cells were maintained in DMEM with 4.5 g/L glucose and L-glutamine and 10% FBS at 37°C and 5% CO_2_. Conditional Gps2^flox/flox^ mice were generated by inGenious Targeting Laboratory. Wild-type mice used as control for all analyses presented here were littermates Gps2^flox/flox/^CD19Cre^−^.

MDA-MB-231 breast cancer cells were grown in DMEM with 4.5 g/L glucose and L-glutamine and 10% FBS at 37°C and 5% CO_2_. MB231 GPS2 KO cells were generated and described previously^47^.

### Protein extraction, subcellular fractionation, immunoprecipitation and western blotting

For whole cell extraction, cultured cells were pelleted and incubated for 20 minutes on ice in IPH buffer (50 mM Tris HCl [pH 8.0], 250 mM NaCl, 5 mM EDTA, 0.5% NP-40, 0.1mM PMSF, 2mM Na3VO4, 50mM NaF, and 1X protease inhibitors (Sigma Aldrich)). For cytoplasmatic, mitochondrial and nuclear extracts fractionation, cells were pelleted and resuspended in gradient buffer (10 mM HEPES [pH 7.9], 1mM EDTA, 210 mM Mannitol, 70mM Sucrose, 10mM NEM, 50 mM NaF, 2 mM Na2VO3, 1mM PMSF and 1x protease inhibitor mixture), then homogenized by syringe followed by low-speed centrifugation for 10 min. The supernatant containing cytosolic proteins was recovered and subjected to high-speed centrifugation to separate the mitochondrial pellet from the cytoplasmic fraction, and the nuclear pellet was lysed for 20 min in high-salt buffer (20 mM Tris-HCL [pH 8.0], 25% glycerol, 420 mM NaCl, 1.5 mM MgCl2, 0.2 mM EDTA, 0.5 mM DTT, 10mM NEM, 50 mM NaF, 2 mM Na3VO4, 1mM PMSF and protease inhibitor mixture). The mitochondrial pellet was incubated for 15 min in lysis buffer (50 mM Tris/HCl [pH 8.0], 300 mM NaCl, 1mM EDTA, 1% Triton X-100, 10mM NEM, 50 mM NaF, 2 mM Na2VO3, 1mM PMSF and protease inhibitor mixture). The concentration of protein extracts was measured using the Bradford assay (Bio-Rad). Extracts were boiled in SDS sample buffer and loaded directly on precast Bio-Rad gels. For immunoprecipitation, protein extracts were incubated with the specific antibody overnight at 4 °C after adjusting the buffer to a final concentration of 150 mM NaCl and 0.5% NP-40 and then incubated for 1 h with Protein A-Sepharose™ 4B (Invitrogen), washed extensively, separated by electrophoresis, transferred onto PVDF membranes (Millipore), and subjected to Western blotting following standard protocols.

### Recombinant Protein Expression

cDNAs encoding the tandem ubiquitin-binding entity (Vx3K0 and Rx3(A7)) (Sims J., 2012, Nature protocol) were subcloned into pET28a-His-tag expression vector. Human clone of GPS2 and mouse clone of PABPC1 and mutated PABPC1 (with four lysine mutations: K78R, K188R, K284R and K512R) were subcloned into pET-32a-His-tag expression vectors. His-tagged Vx3K0, Rx3(A7), GPS2 and PABPC1 were produced as a His-tag fusion protein in BL21 *Escherichia coli*, resin-purified on nickel-nitrilotriacetic acid beads, and eluted accordingly to manufacturer’s protocol (Life Technologies). The His-Vx3K0/Rx3(A7)-conjugated agarose were stored at 4°C in PBS supplemented with 30% glycine.

### Immunochemical methods

For pull-down of K63-ubiquitylated proteins, cells were lysed in high-stringency buffer (50 mM Tris, pH 7.5; 500 mM NaCl; 5 mM EDTA; 1% NP40; 1 mM dithiothreitol (DTT); 0.1% SDS) containing 1.25 mg ml−1 N-ethylmaleimide, 0.1% PMSF, and protease inhibitor cocktail (Roche). His-Vx3K0/Rx3(A7) were added to immobilize the K63-ubiquitylated proteins, and bound material was washed extensively in high-stringency buffer four times. Proteins were resolved by SDS-APGE and analyzed by immunoblotting.

### In vitro ubiquitination assay

Ubiquitination assays were carried out at 30°C for 2h in 50mM Tris-HCl, pH7.6, containing 50mM NaCl, 5mM MgCl2, 5mM ATP, 1x Ubiquitin Aldheide, 50nM E1, 5μg ubiquitin, 200 nM UbcH13– Uev1a E2 complexes (Boston Biochem), 0.1ug recombinant human MUL1 Protein (H00079594-P01, Novus Biologicals). Bacterial expressed His-PABPC1/mutated PABPC1 was used as substrate.

### Mitochondrial RNA Isolation and RNA-seq

Mitochondrial RNA was isolated from isolated mitochondrial pellet following the manufacturer protocol for the RNeasy Kit (QIAGEN). Isolated mitochondrial RNA was subjected to quality control on Agilent Bioanalyzer and RNA library preparation following Illumina’s RNA-seq Sample Preparation Protocol. Resulting cDNA libraries were sequenced on the Illumina’s HiSeq 2000.

### Mitochondrial Content

Total DNA was extracted from cells using QuickExtract DNA Extraction Solution 1.0 (Epicenter) following manufacturer’s instructions. DNA amplification of the mitochondrial-encoded NADH dehydrogenase 1 (mt-ND1) relative to nuclear TFAM was used to determine mitochondrial DNA copy number.

#### Translation activity

Puromycin incorporation into newly synthesized proteins was performed to measure translation activity. MEFs WT and GPS2-KO cells were treated in the presence or absence of homoharringtonine (HHT) (5 μM, Tocris Bioscience) for 10 min to prevent new translation initiation, prior to an additional 5 min incubation with puromycin (100 μM, Sigma) and emetine (200 μM, Sigma) at 37°C. Cells were collected by centrifugation, and lysate was prepared as described above. Thirty micrograms of protein were loaded onto a 10% SDS-PAGE gel for immunoblotting analysis.

### Sample preparation for MS analysis

For mitochondrial proteome profiling, mitochondrial proteins from MEFs WT and GPS2-KO cells were isolated as described before^9^. An equal amount of solubilized mitochondrial proteins from different samples were separated by SDS-PAGE and then digested in-gel by trypsin overnight using standard methods^74^.

For SILAC-K63 experiments, MEFs WT and GPS2-KO cells or MDA-MB231 WT and GPS2-KO cells were grown in medium containing native (unlabeled) L-arginine and L-lysine (Arg0/Lys0) as the light condition, or stable isotope-labelled variants of L-arginine and L-lysine (Arg10/Lys8) as the heavy condition^75^. Proteins from whole cellular lysates were extracted from SILAC-labelled MEFs WT and GPS2-KO cells or MDA-MB-231WT and GPS2-KO cells as descripted before. An equal amount of proteins from the two SILAC states was mixed and precipitated by His-Vx3K0-conjugated agarose and incubating at 4°C for 1h. Precipitated proteins were eluted with SDS sample buffer, incubated with 10 mM DTT for 10min at 100 °C. Proteins were separated by SDS-PAGE and then digested in-gel by trypsin overnight using standard methods. The resulting peptides were desalted using reverse phase (C18) Tips (Thermo Scientific) per the manufacturer’s instructions.

For TMT based-K63 experiments, MEFs WT and GPS2-KO cells were grown in normal medium to 80% confluency. After lysis with IPH buffer, protein quantities from whole cellular lysates were determined using the Bradford assay. An equal amount of proteins from MEFs WT or GPS2-KO cells was precipitated by His-Vx3K0-conjugated agarose and incubating at 4°C for 1h. Precipitated proteins were eluted with SDS sample buffer, incubated with 10 mM DTT for 10min at 100 °C. Proteins were separated by SDS-PAGE and then digested in-gel by trypsin overnight using standard methods. The resulting peptides were desalted using C18 Tips (Thermo Scientific) per the manufacturer’s instructions and vacuum-dried. The dried peptides were redissolved in 0.5 M TEAB, and processed independently according to the manufacturer’s protocol for a 10-plex Tandem Mass Tag (TMT) reagent labeling kit (Thermo Fisher Scientific). The different TMT-labeled peptide mixtures were pooled equally, desalted and dried by vacuum centrifugation.

### Mass spectrometric analysis

Peptides were analyzed on a Q-Exactive HF mass spectrometer (QE-HF, Thermo Fisher Scientific) equipped with a nanoflow EasyLC1200 HPLC system (Thermo Fisher Scientific). Peptides were loaded onto a C18 trap column (3 μm, 75 μm × 2 cm, Thermo Fisher Scientific) connected in-line to a C18 analytical column (2 μm, 75 μm × 50 cm, Thermo EasySpray) using the Thermo EasyLC 1200 system with the column oven set to 55 °C. The nanoflow gradient consisted of buffer A (composed of 2% (v/v) ACN with 0.1% formic acid) and buffer B (consisting of 80% (v/v) ACN with 0.1% formic acid). For protein analysis, nLC was performed for 180 min at a flow rate of 250 nL/min, with a gradient of 2-8% B for 5 min, followed by a 8-20% B for 96 min, a 20-35% gradient for 56min, and a 35-98% B gradient for 3 min, 98% buffer B for 3 min, 100-0% gradient of B for 3 min, and finishing with 5% B for 14 min. Peptides were directly ionized using a nanospray ion source and analyzed on the Q-Exactive HF mass spectrometer (Thermo Fisher Scientific).

QE-HF was run using data dependent MS2 scan mode, with the top 10 most intense precursor ions acquired per profile mode full-scan precursor mass spectrum subject to HCD fragmentation. Full MS spectra were collected at a resolution of 120,000 with an AGC of 3e6 or maximum injection time of 60 ms and a scan range of 350 to 1650 m/z, while the MS2 scans were performed at 45,000 resolution, with an ion-packet setting of 2e4 for AGC, maximum injection time of 90 ms, and using 33% NCE. Source ionization parameters were optimized with the spray voltage at 2.1 kV, transfer temperature at 275 °C. Dynamic exclusion was set to 40 seconds.

### Data analysis

All acquired MS/MS spectra were searched against the Uniprot mouse complete proteome FASTA database released on 2013_07_01, using the MaxQuant software (Version 1.6.7.0) that integrates the Andromeda search engine. Enzyme specificity was set to trypsin and up to two missed cleavages were allowed. Cysteine carbamidomethylation was specified as a fixed modification. Methionine oxidation, N-terminal acetylation, and lysine ubiquitination were included as variable modifications. Peptide precursor ions were searched with a maximum mass deviation of 6 ppm and fragment ions with a maximum mass deviation of 20 ppm. Peptide and protein identifications were filtered at 1% FDR using the target-decoy database search strategy. Proteins that could not be differentiated based on MS/MS spectra alone were grouped to protein groups (default MaxQuant settings). For mitochondrial protein annotation, we utilized the latest Mitocarta dataset(https://www.broadinstitute.org/files/shared/metabolism/mitocarta/human.mitocarta3.0.html).

### Statistical analysis

The statistical data are from three independent experiments. Results are shown as mean ± SEM unless mentioned otherwise. Statistical analysis was performed by the Student’s t-test for TMT-based quantification and RNA-seq data.

---

**Supplemental Figure 1, Related to Figure 1**

A, Western blot analysis of GPS2 expression in GPS2 WT and KO MEFs cells. Anti-a-Tub blot was used as loading control.

B, Mitochondrial DNA content in WT and GPS2 KO MEFs cells. **indicate p-value<0.01.

C, Gene ontology analysis of 891 differentially expressed proteins induced by GPS2-deficiency from GPS2 WT and KO MEFs.

**Supplemental Figure 2, Related to Figure 2**

A, Comparison of the binding affinity of Vx3K0 and Rx3 to different type and length ubiquitin chains.

B, WB analysis of K63 ubiquitinated proteins captured by Vx3K0 from whole cell extracts in GPS2 WT and KO MEFs.

C, Scatter plotting analysis of SILAC data obtained from two independent SILAC-K63 ubiquitome experiments. The KO/WT SILAC ratio of two labeled samples were converted to log2 scale for analysis. The proteins identified by SILAC-K63 ubiquitome and their SILAC ratios are listed in supplemental Table 3.

D, TMT labeling-based K63 proteomic profiling of GPS2 WT and KO MEFs. Each sample (WT/KO) has 3 replicates for LC-MS/MS analysis.

E, Gene ontology analysis of 22 putative K63 ubiquitinated proteins whose ubiquitination are mediated by GPS2 identified in MB231.

**Supplemental Figure 3, Related to Figure 2**

A, Venn diagram comparing identified putative K63 ubiquitinated proteins whose ubiquitination are mediated by GPS2 from MEFs SILAC, MEFs TmT and MDA-MB231 SILAC datasets.

B, Detection of K63 ubiquitination of eif3M purified by specific K63-ubiquitin binding domain Vx3K0 by IP/WB using fractionated mitochondrial extracts in GPS2 WT, KO and KO knockdown Mulan (siMul1) MEFs.

