## Supplementary figures and images for "Inhibition of Mul1-mediated ubiquitination promotes mitochondria-associated translation"

### Supp Figure 1

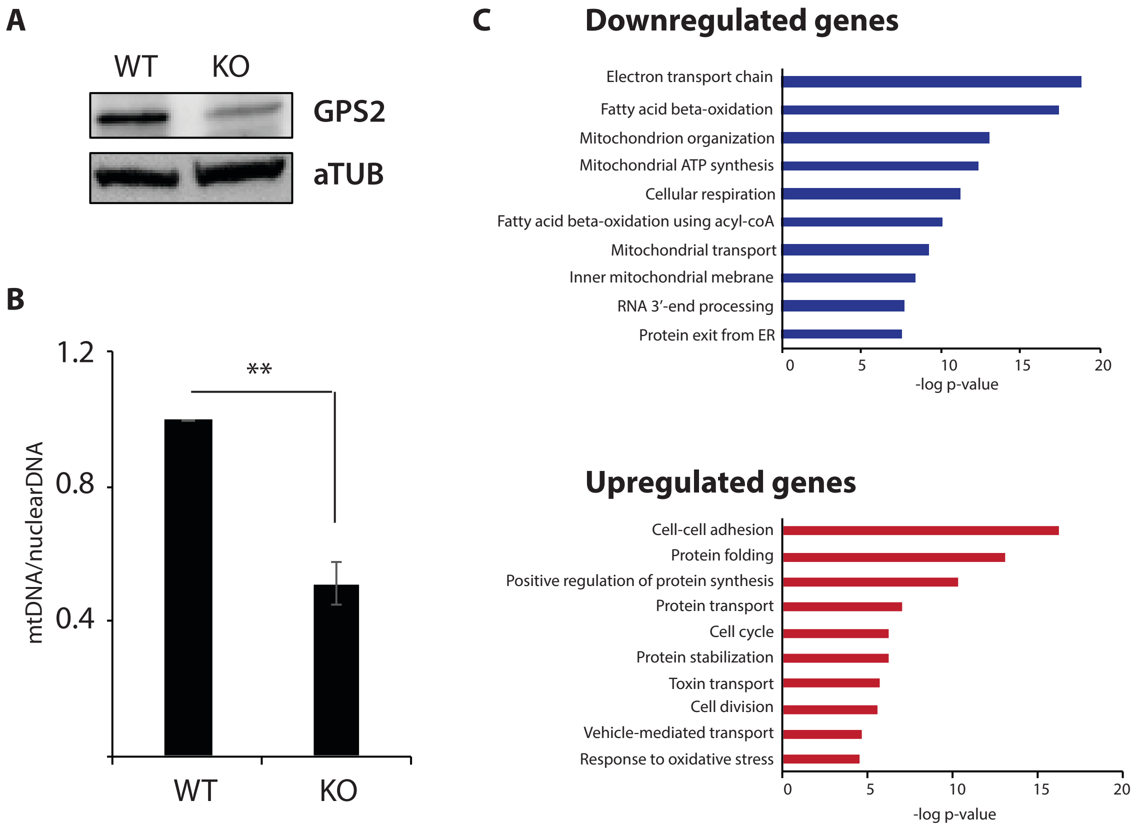


**Supp Figure 1**

**Gao et al., 2021**

### Supp Figure 2

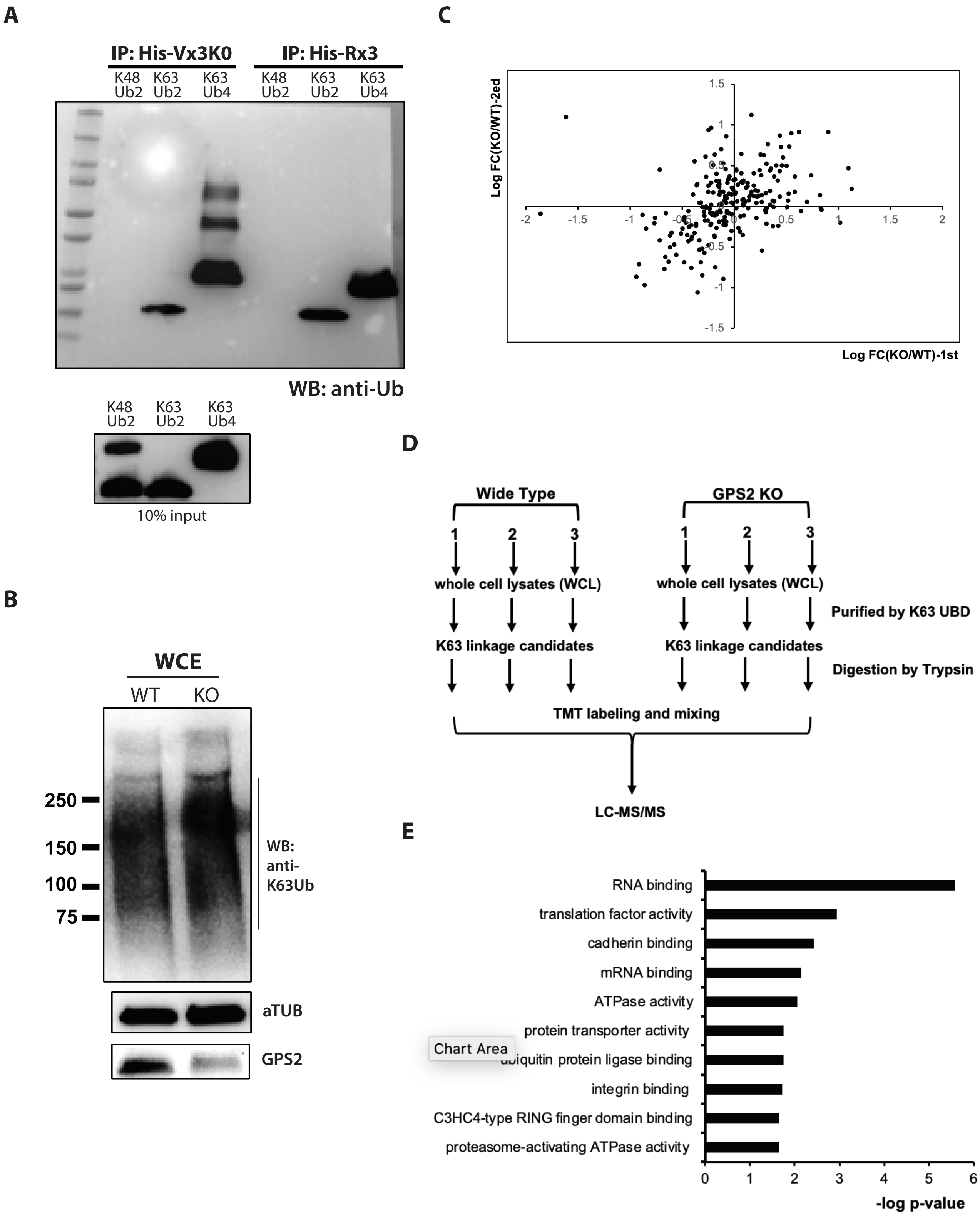


Supp Figure 2

Gao et al., 2021

### Supp Figure 3

**A**

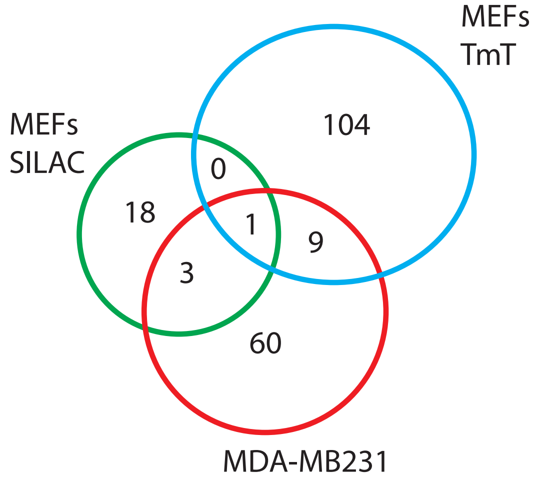


**B**

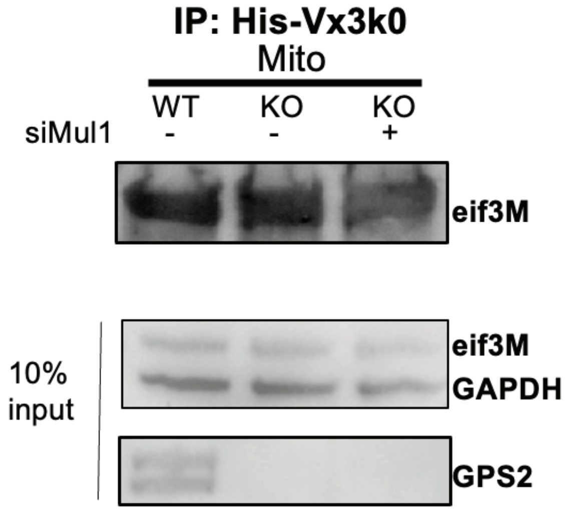


Supp Figure 3
Gao et al., 2021
